# Sequence composition and conservation predict the phenotypic relevance of transposable elements

**DOI:** 10.1101/2024.03.11.584453

**Authors:** Yari Cerruti, Daniela Fusco, Mahdis Jahanbin, Davide Marnetto, Paolo Provero

## Abstract

Transposable elements (TEs) are powerful drivers of genome evolution, in part through their ability to rapidly rewire the host regulatory network by creating transcription factor binding sites that can potentially be turned to the host’s advantage in a process called exaptation. However, the effects on the host phenotype vary widely among different TEs. Here, we classify TEs based on their contribution to the human host phenotypes at both molecular and macroscopic scales. TE contributions to chromatin accessibility, gene expression, and the heritability of complex traits are strongly correlated to each other, confirming that the main mechanism through which TEs affect the host phenotype is through the rewiring of the regulatory network. TE sequence and evolutionary features are able to explain a large fraction of the variance in phenotypic relevance, and in particular more than 50% of the variance of their contribution to the heritability of complex traits. A conspicuous exception to this pattern is represented by TEs of the ERV1 family, whose phenotypic impact cannot be explained by our model: In particular, this family includes a set of relatively young TEs whose phenotypic relevance is much larger than would be expected based on their sequence and evolutionary parameters. These TEs are involved in fast-evolving biological processes related to the interaction of the organism with its environment. In conclusion, our results confirm quantitatively that TE insertions affect the host phenotype mostly through the rewiring of its regulatory network; identify a signature of phenotypic relevance based on sequence and conservation properties; and highlight several TEs as promising candidates for functional studies.

## Introduction

Transposable elements (TEs) have been shown to provide raw material for the rapid evolution of genomes, and specifically of gene regulation, by creating quickly dispersing genetic elements potentially exploitable by the host as binding sites for factors involved in regulating transcription and three-dimensional genome conformation (see [1–3] for recent reviews), in a phenomenon called TE exaptation. Human regulatory network rewiring by TE exaptation has been recently shown to be relevant to a variety of biological processes, including immune response [4–7], germ cell development [8,9], longevity [10], pluripotency [11], and 3D genome conformation [12].

TE exaptation can be used by the host not only to rewire the regulatory network by providing new regulatory targets of transcription factors (TFs), but also to increase the robustness of the existing regulatory network by creating additional, redundant binding sites near existing ones [13–15]. This fact suggests that methods based on the analysis of TE sequence and the selective pressure acting on it might usefully complement those based on genetic perturbations, as the latter might be prone to false negatives when the role of the exapted TEs is to provide redundancy. Analytical approaches based on sequence properties and evolutionary conservation have the added advantage of bypassing the need to identify the tissue, cell type, or biological condition in which the role of the exapted TEs is exerted.

In this work we first classify TEs based on their phenotypic effects on the host at both the molecular and the macroscopic levels, and investigate the relationships between these levels. At the molecular level, we consider chromatin accessibility and gene expression as phenotypes, while at the macroscopic level we investigate the heritability of complex traits. We then identify sequence and conservation features that are predictive of such effects.

## Results

### Macroscopic phenotypic effects of TEs are driven by regulatory intermediate molecular phenotypes

We considered three measures of phenotypic relevance that can be associated with each TE. Two of them refer to molecular phenotypes (chromatin accessibility and gene expression), while one refers to macroscopic traits, namely complex traits as measured in genome-wide association studies (GWASs).

Specifically (see Methods for details) we considered, for each TE, its effect on

- *Chromatin accessibility*, evaluated as the fraction of TE sequence in the genome that overlaps an ENCODE candidate cis-reguatory element (cCRE).
- *Gene expression*, evaluated as the number of genes with at least one expression quantitative trait locus (eQTL) residing inside a copy of the TE in at least one GTEx tissue, divided by the number of common variants found in the TE.
- *Complex trait heritability*, evaluated as the heritability enrichment as computed in [16] through a meta-analysis of dozens of GWAS studies.

We first asked to what degree these measures of phenotypic relevance agree with each other. As shown in Fig. 1A, all pairwise correlations between the three measures are positive and highly significant (all *P* < 2.2 * 10^-16^). As expected, the two molecular phenotypes show high concordance (Spearman’s ρ = 0.59), confirming that the effect of TE exaptation on gene expression is largely driven by the creation of regulatory elements.

**Fig. 1.**
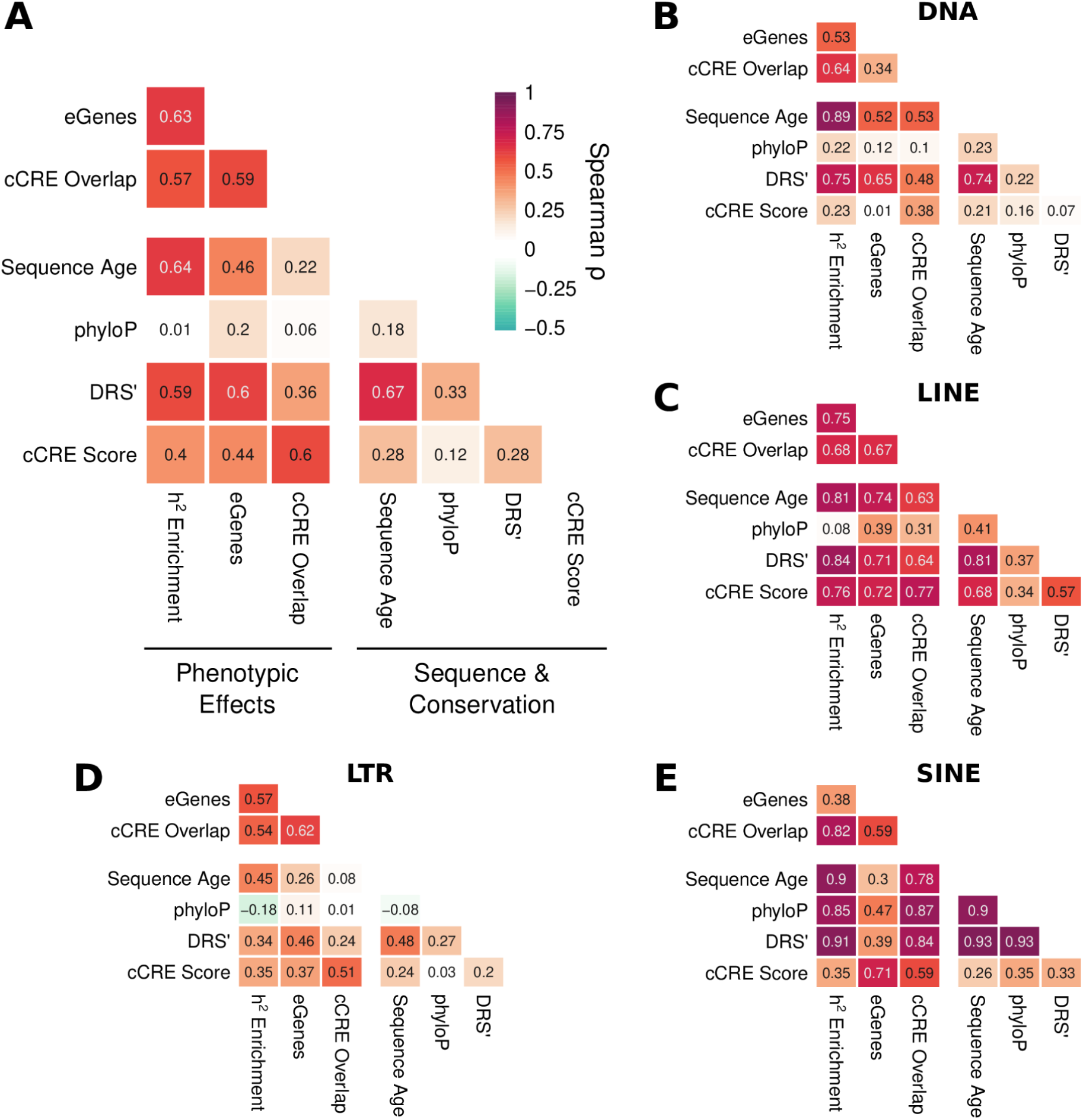
A: Spearman correlation between measures of phenotypic impact and sequence/conservation features. B-E: Same by TE class.

Perhaps less obvious is the high correlation between effects at the molecular and macroscopic levels (ρ = 0.57 and 0.63 of complex trait heritability with chromatin accessibility and gene expression, respectively). These results generally apply to all TE families (Fig. 1B-E), and suggest that the contribution of TEs to complex trait variability is largely driven by their effects on the regulatory network. Macroscopic traits are the targets of selection, and complex traits in particular are known to be subjected to stabilizing selection [17]. We therefore followed up by assessing whether TEs with consensus (i.e. ancestral) sequence suitable to assume a regulatory role in the human host would be subjected to purifying selective pressure.

### Consensus sequence composition predicts evolutionary conservation and contemporary selective pressure

To classify TEs based on their ability to create regulatory elements, we used gkmSVM [18] to train a support vector machine (SVM) model able to predict the cCRE status of a sequence from its k-mer composition (see Methods). To avoid circularity, the model was trained on ENCODE cCREs not overlapping TEs, and, in cross-validation, achieved good predictive performance (area under the ROC curve [AUC] = 0.793). The model was then applied to the consensus sequence of each TE (derived from the Dfam database [19], with some modifications, see Methods). The score assigned by gkmSVM to each TE, henceforth referred to as the cCRE score, showed good correlation with the fraction of TE sequence overlapping ENCODE cCREs (Spearman ρ = 0.60, *P* < 2.2 * 10^-16^), confirming that the sequence determinants of chromatin accessibility are similar for TE-derived and non-TE-derived genomic regions. Having been obtained from the consensus, i.e. ancestral, sequence, this result confirms that for most TEs the regulatory potential is present immediately upon insertion in the genome, as argued in [13].

To investigate the relationship between cCRE score and selection, we evaluated three measures of purifying selective pressure on each TE (see Methods for details):

- Sequence age, namely the oldest human ancestor in which the sequence was incorporated in the genome, as determined in [20]. For each TE we considered the median age across all copies. This evaluation of TE sequence age was in excellent agreement with that derived in [16] based on the divergence of individual copies from the consensus sequence (see Suppl. Fig. 1).
- phyloP score [21] based on the alignment of 241 mammalian genomes [22], which measures the degree of conservation of each TE copy among the mammals that share it. For each TE we considered the average phyloP score over all bases covered.
- The depletion rank score (DRS) [23], measuring the depletion of variants in a modern population. Also this measure was averaged over all bases covered by each TE class. Throughout the paper we use DRS’, defined as 1-DRS/100, so that higher values of DRS’ correspond to greater conservation (greater depletion of variants) in agreement with the other two measures.

These three measures can be considered as representing purifying selective pressure at different time scales: Sequence age refers to the oldest ancestor that can be still inferred as having carried each specific copy of a TE; the phyloP score measures the degree to which the sequence of such copies has been conserved; while DRS’ measures the degree of purifying selection acting on each genomic region in modern human populations.

As expected, the three measures were positively correlated with each other (Fig. 1A). Perhaps surprisingly, the strongest correlation was observed between DRS’ and sequence age, showing that, at least in the case of TEs, the oldest sequences are the ones currently experiencing the highest degree of purifying selective pressure.

Selective pressure was moderately and positively associated with the cCRE score (ρ of the selection measures between 0.12 and 0.28). Stratifying by TE family we observe that such correlation is driven in particular by LINE elements (Fig. 1C). Thus the TEs that have been conserved during mammalian evolution and are still subject to selective pressure in modern humans tend to be those whose sequence is more favorable to an open chromatin state.

### Sequence composition and conservation patterns explain the variance of phenotypic relevance among TE classes

Having defined these measures of sequence composition and conservation, and assessed their mutual relationships, we asked to what extent they were predictive of the molecular and macroscopic phenotypic effects of TE insertions defined above. We first evaluated separately the cCRE score and the three measures of conservation as predictors of phenotypic relevance, using univariable linear regression. This revealed that (Tab. 1, Suppl. Tab. 1-2):

- phyloP scores were consistently the weakest predictors of every measure of phenotypic relevance.
- As expected, the cCRE score was the strongest predictor of chromatin accessibility, explaining by itself ∼31% of its variance among TEs.
- DRS’ was the best predictor of the effect on gene expression, explaining ∼30% of its variance.
- Finally, sequence age was the strongest predictor of enrichment in complex trait heritability, as previously suggested [16], explaining ∼44% of its variance.

While the four predictors of phenotypic relevance are correlated to each other (see Fig. 1A), their variance inflation factors are all below 2, so that a multivariable regression model using them as multiple regressors can be interpreted without problems arising from multicollinearity. We thus fitted such a multivariable model to predict each of the three measures of phenotypic relevance, using the cCRE score and the three measures of selection as regressors, and including all the pairwise interactions between them. The models achieved high fractions of variance explained, with adjusted *R^2^*values ranging from 0.44 for gene expression to 0.58 for complex trait heritability. Tab. 1 shows coefficients and *P*-values for the significant predictors of complex trait heritability. The strongly significant interaction terms between selection measures suggest a complex interplay of selection patterns at different time scales in predicting complex trait heritability. The corresponding results for the gene expression and chromatin accessibility are shown in Suppl. Tab. 1 and Suppl. Tab. 2.

**Tab. 1.** Regression coefficients and P-values of linear models predicting complex trait heritability enrichment from sequence features and measures of conservation. The adjusted R^2^ of the multivariable model is 0.58. UNI: univariable model. MULTI: multivariable model including all pairwise interactions.

| | $\beta_{\text{UNI}}$ | $P_{\text{UNI}}$ | $\beta_{\text{MULTI}}$ | $P_{\text{MULTI}}$ |
| --- | --- | --- | --- | --- |
| **sequence age** | 0.6665 | $3.889 \cdot 10^{-102}$ | 0.3959 | $5.006 \cdot 10^{-26}$ |
| **DRS'** | 0.489 | $1.683 \cdot 10^{-48}$ | 0.3945 | $6.111 \cdot 10^{-19}$ |
| **phyloP** | 0.3115 | $3.745 \cdot 10^{-19}$ | -0.4966 | $5.967 \cdot 10^{-14}$ |
| **cCRE score** | 0.3214 | $2.342 \cdot 10^{-20}$ | 0.1921 | $5.19110^{-12}$ |
| **sequence age:DRS'** | | | 0.1704 | $9.327 \cdot 10^{-12}$ |
| **sequence age:phyloP** | | | 0.2137 | $1.299 \cdot 10^{-6}$ |
| **phyloP:DRS'** | | | 0.2314 | $2.344 \cdot 10^{-6}$ |
| **cCRE score:phyloP** |  |  | 0.08076 | 0.01888 |
| **cCRE score:DRS'** |  |  | -0.07438 | 0.03166 |
| **cCRE score:sequence age** |  |  | -0.0619 | 0.1188 |

#### Older TEs of all classes contribute to complex trait heritability by rewiring developmental regulatory networks

Fig. 2A shows the prediction of the model for heritability enrichment vs. its actual (inverse normal transformed) value. The top 10 TEs by predicted heritability enrichment are labelled in the figure and listed in Tab. 2. In agreement with the dominant role played by sequence age in predicting heritability enrichment, these are all older TEs whose origin can be traced to shortly after the origin of mammals. We used GREAT [24] to investigate the functional characterization (Gene Ontology Biological Process, GO:BP) of these TEs, by merging together the respective genomic regions. The top GO:BP terms, after redundancy reduction [25], are represented in Fig. 2B and are largely dominated by developmental processes. Complete enrichment results are reported in Suppl. Data 1, while the results of enrichment analyses for the individual TEs are reported in Suppl. Data 2. Beside cases already examined in the literature (see Discussion), we highlight two novel functional associations with members of the Tigger family, namely, Tigger14a and Tigger16b, that show their strongest enrichment in developmental processes related, respectively, to pigmentation and the eye (see Suppl. Data 2).

**Fig. 2:**
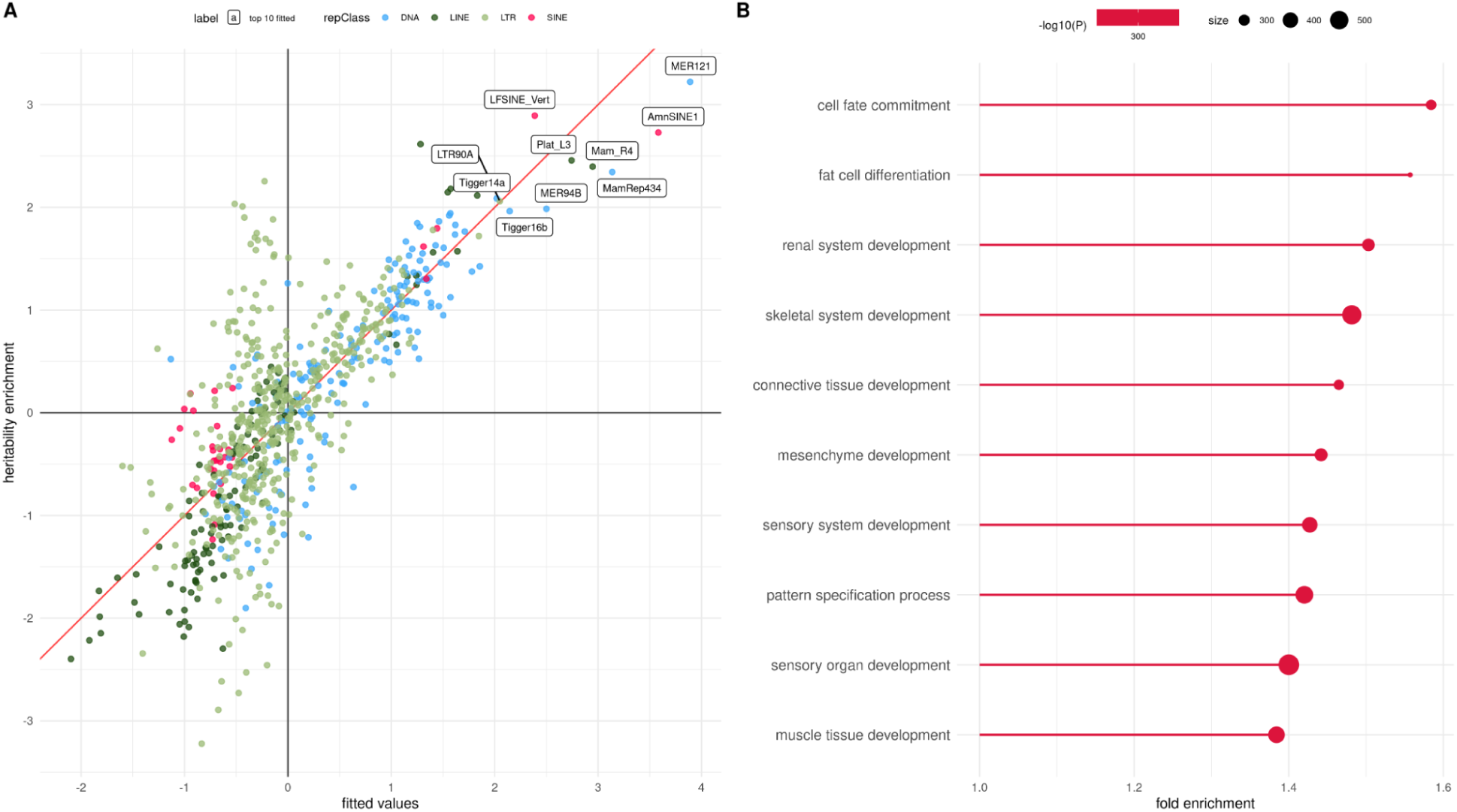
A: Linear model predictions (x axis) vs actual heritability enrichments (after inverse normal transformation) of TEs. The labels indicate the top 10 TEs by predicted value. B: Gene Ontology terms enriched in top 10 TEs by predicted value.

**Tab. 2.**
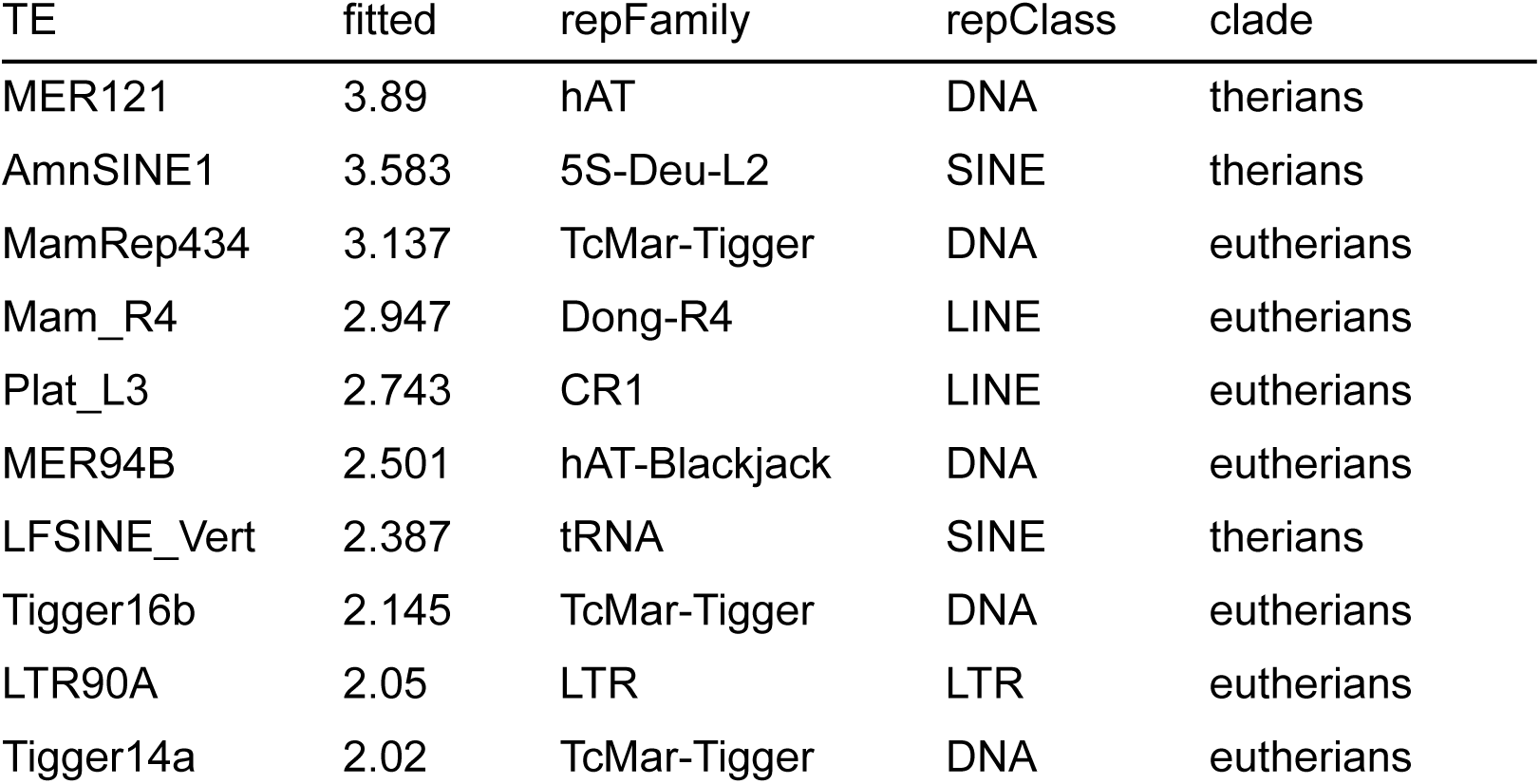
The 10 top TEs in terms of predicted complex trait heritability enrichment (“fitted” column). The clade of origin corresponds to the median sequence age of the TE copies (Methods)

#### Younger LTRs disproportionately contribute to complex trait heritability by rewiring regulatory networks involved in the interaction with the environment

As discussed above, the model we developed explains a large fraction of the heritability enrichment. However, Fig. 2A clearly shows a large group of TEs of the LTR class with heritability enrichment higher than what is predicted by the model. In Fig. 3A, where LTR families are highlighted, it is evident how LTRs specifically belonging to the ERV1 family show a behavior that deviates from the model predictions. The top 10 TEs by model residual (labelled in Fig. 3A) are shown in Tab. 3. These results suggest the existence of a set of LTRs, mostly belonging to the ERV1 family and whose insertion can be traced to after the origin of primates, whose effect on heritable traits is much higher than can be predicted from their sequence and evolutionary properties.

**Fig. 3.**
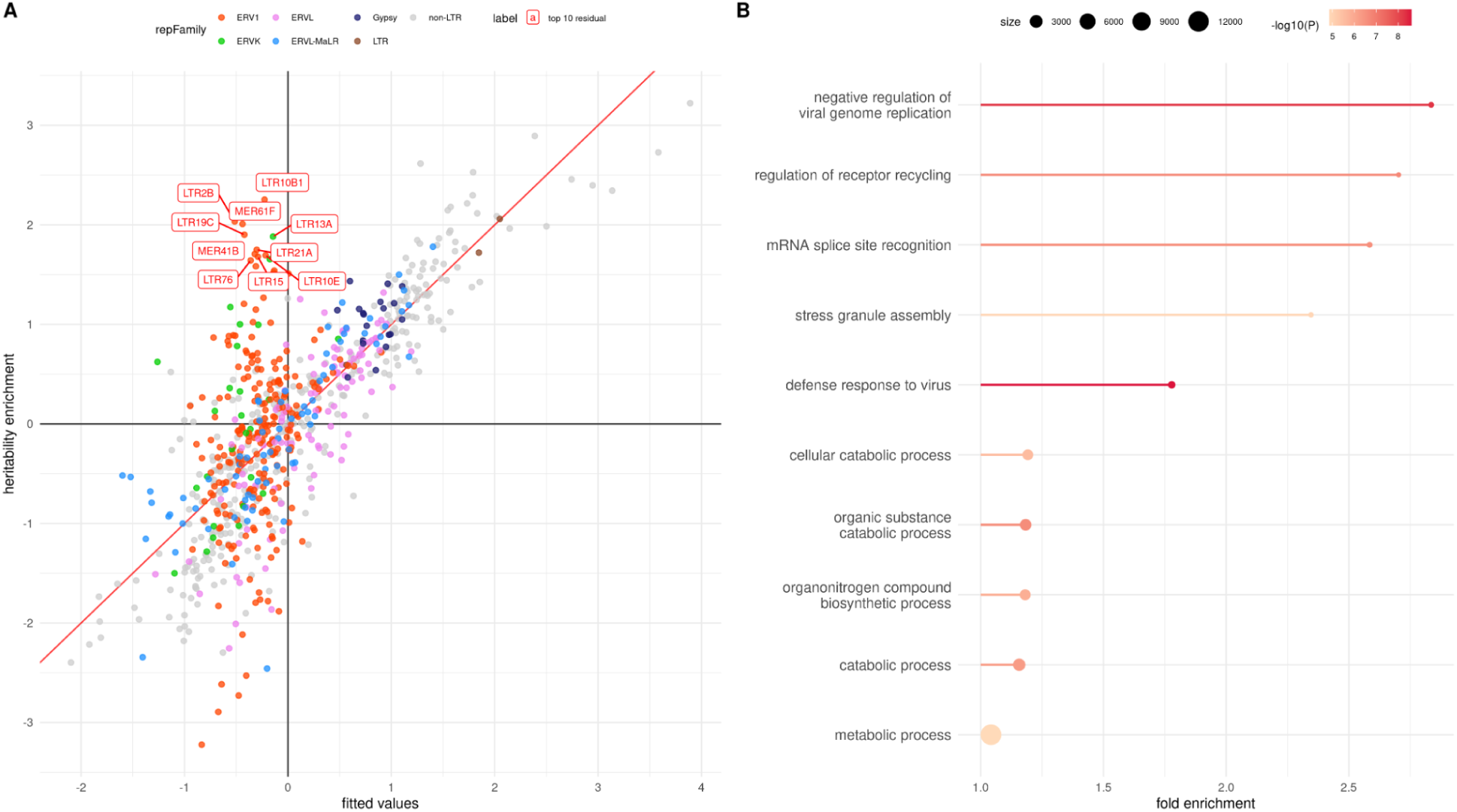
A: Same as Fig. 2A, except that the labels now indicate the top 10 TEs by model residual. B: Gene Ontology terms enriched in the top 10 TEs by model residual.

**Tab. 3:** The 10 top TEs in terms of model residual.

| TE | residual | repFamily | repClass | clade |
| --- | --- | --- | --- | --- |
| LTR2B | 2.548 | ERV1 | LTR | apes |
| LTR10B1 | 2.48 | ERV1 | LTR | catarrhini |
| MER61F | 2.448 | ERV1 | LTR | simians |
| LTR19C | 2.323 | ERV1 | LTR | catarrhini |
| LTR21A | 2.051 | ERV1 | LTR | catarrhini |
| MER41B | 2.027 | ERV1 | LTR | simians |
| LTR13A | 2.026 | ERVK | LTR | catarrhini |
| LTR76 | 2.003 | ERV1 | LTR | catarrhini |
| LTR15 | 1.971 | ERV1 | LTR | catarrhini |
| LTR10E | 1.91 | ERV1 | LTR | catarrhini |

The GO:BP enrichments of these TEs, considered together, were less prominent, in terms of both number of significant terms and *P*-values, than those found for the top predicted heritability enrichment TEs. The most significant GO terms are shown in Fig. 3B, and complete enrichment results are found in Suppl. Data 3. Many of these GO:BP terms refer to fast-evolving traits related to the organism/environment interaction, such as immune response and metabolism. Among the strongest GO:BP enrichments of these individual TEs (Suppl. Data 4), we mention that LTR76 appears to be associated with many virus- and interferon-related processes, LTR19C to inflammation, and MER61F to the metabolism of uronic acid.

Hence, the model we developed to predict heritability enrichment from sequence composition and conservation patterns does not apply to the ERV1 family of TEs. Indeed, while the model with all TEs explained 58% of the variance in complex trait heritability enrichment, the same model restricted to ERV1 TEs predicted just 13% of the same variance; conversely, when the ERV1 family was removed from the dataset, the fraction of variance explained increased to 75%.

Analyzing top-predicted and top-residual TEs for the molecular measures of phenotypic relevance (gene expression and open chromatin) we obtained qualitatively similar results, shown in Suppl. Tabs. 3-6, Suppl. Figs. 2-5, and Suppl. Data 5-8.

## Discussion

We have shown that sequence and evolutionary features of TEs predict their phenotypic relevance at the macroscopic and molecular level. Our results confirm quantitatively that the effects of TEs on the human host’s phenotype are largely driven by their exaptation as regulatory elements and the ensuing changes in gene expression. The most important predictors are sequence age and the density of variants in modern populations, representing the opposite ends of the selection time scale. The TEs with the greatest impact on human phenotype are thus those that inserted themselves in the genome shortly after the origin of mammals, and are depleted in genetic variation in modern humans. Enrichment analysis shows these TEs to be mostly involved in the regulation of developmental pathways.

Among these TEs, MER121 is particularly intriguing, since it was identified in [26] as an example of a repeated element subject to extreme selective pressure, although, to the best of our knowledge, its function has not been further elucidated. Enrichment analysis (see Suppl. Data 2) suggests its involvement in neurogenesis and the morphogenesis of several organs including heart, kidney, and lung. MamRep434 and LFSINE_Vert were recently suggested [27] to be involved in gene regulation of glutamatergic neuron precursors, and indeed our enrichment analysis identifies “generation of neurons” and “synaptic transmission, glutamatergic” as significantly enriched for both TEs (Suppl. Data 2). AmnSINE1 has also been found to provide regulatory elements involved in the development of the mammalian brain (see e.g. [28]), again in agreement with the enrichment results, but also to be involved in innate immunity [4,7]. Other TEs, such as Tigger14a and Tigger16b have not been associated with specific functions in the literature, and thus would be interesting targets of experimental investigation.

In addition, we have shown that some LTRs do not follow the general pattern of association between sequence/conservation features and phenotypic relevance. Most of these belong to the ERV1 family, and include some TEs whose phenotypic impact, in particular on the heritability of complex traits, is much larger than expected from their sequence and conservation features. These TEs integrated themselves in the genome after the origin of primates, and are functionally related to fast-evolving pathways involved in the interaction with the environment, such as response to pathogens and metabolism. Intriguingly, two of these TEs (LTR2B, and MER41B) have been associated [29] to trophoblast gene expression. The placenta is indeed among the fastest-evolving organs in mammals [30], a fact which is compatible with the involvement of relatively young TEs. Moreover, LTR10E and LTR10B1 have been implicated in the rewiring of the TP53 regulome in primates [31].

The main limitation of our study is the fact that all the measures we considered were averaged or otherwise summarized over all the copies of a TE: Future investigations should consider the differences in sequence and conservation features among the different copies of a TE to fine-map the individual copies that were actually exapted by the host and their regulatory function. Similarly, our measures of phenotypic relevance are summarized across traits (for heritability), or tissues/cell lines (for gene expression and chromatin): Analyzing these biological contexts separately could provide further insight on TE relevance. Finally, we have not directly investigated the role of TE transcription, which is known to be relevant beyond their regulatory role especially in the very early stages of development, such as zygotic genome activation (see [32] for a recent review).

In conclusion, we have shown that TE sequence and conservation features are strongly predictive of their functional impact on the host phenotype through rewiring of the gene regulatory network.

## Methods

### TE data preprocessing

Bed files containing genome-wide coordinates of repetitive elements were retrieved from the latest UCSC Repeatmasker [33] GRCh38 annotation, and only transposable elements (repeatmasker repClass annotation “LINE, “SINE”, “LTR”, “DNA”, or “Retroposon”) were retained. TEs mapping on X, Y, and mitochondrial chromosomes were excluded from further analysis. TEs mapping within the Human Leukocyte Antigen (HLA) region of chromosome 6 were also excluded, owing to the extreme variability of the HLA region which could skew sequence-based measures of conservation. A total of 906 TEs covering a minimum of 10,000 bases with available PhyloP and DRS values were retained for further analysis. After such a filter the “Retroposon” class contained only 3 TEs, and therefore was not separately analyzed as a class.

### Measures of phenotypic relevance

We chose three measures of phenotypic relevance that could quantify the impact of TEs on molecular and macroscopic traits, which were subsequently used as dependent variables of our linear models. Such measures were computed for each TE (i.e. a repeatmasker “repName”) as described below.

#### Chromatin accessibility

Bed files containing genome-wide annotated functional regions were retrieved from the ENCODE Candidate Cis-Regulatory Elements (cCRE) Genome Browser track [34]. TE coordinates were intersected with cCREs coordinates in order to assess the overlap between TEs and open chromatin regions. The chromatin accessibility of a TE was computed as the number of TE bases overlapping any cCRE divided by the total genomic coverage of the TE.

#### Gene expression

eQTLs coordinates were retrieved for all 49 GTEx tissues [35]. For each TE, we counted the number of eGenes with eQTLs inside the TE, and divided it by the number of common SNPs (MAF > 0.01 in GTEx) inside the TE.

#### Complex trait heritability

The heritability enrichment of TEs was retrieved from Supplementary Data 4 of [16].

### Sequence and Conservation Features

#### Sequence cCRE score

The regulatory content within TEs was predicted from sequence data with a machine learning approach. gkmSVM [18], an SVM machine learning algorithm that is able to predict regulatory sequence features from gapped k-mers, was trained with default parameters on 15,000 randomly chosen cCREs not overlapping any TE. The trained model was then used to classify a set of TE consensus sequences downloaded from the Dfam database [19], resulting in a weight (namely the “cCRE score”) representing the propensity of the TE consensus sequence to function as a cCRE based on its k-mer composition. Since all other predictors were computed for TEs retrieved from repeatmasker, whereas the sequence score was computed on consensus sequences downloaded from Dfam, we could only compute cCRE scores on consensus sequences belonging to 816 TEs, due to differences in nomenclature. For the remaining TEs, the cCRE score was computed by applying the same trained model to the sequence of each copy of each TE, and then averaging the weights over all copies. Supplementary Fig. 6 shows that the two methods highly correlate (R^2^ = 0.88), when tested on 80 random TEs with data available from both sources.

#### Sequence age

A genomic sequence age was assigned to each TE copy by intersecting the TE coordinates with the genomic age estimated in [20] and selecting their “max_age”, which corresponds to the oldest human ancestor in which a copy can be recognized. Since age estimates are available in non-overlapping windows that are highly variable in size (mean = 93.0, sd = 3522.013), TE copies overlapping consecutive windows were assigned the estimated age of the longest fragment. The age of a TE was then computed as the median age across all its copies. This assessment of sequence age, based on multiple alignment of vertebrate genomes, strongly correlates with the one, used e.g. in [16], based on the divergence of copies from the ancestral TE sequence, as shown in Suppl. Fig. 1.

#### phyloP

Genome-wide phyloP scores computed from the multiple alignment of 241 mammalian genomes [22], downloaded from the UCSC Genome Browser, were intersected with TE coordinates using the UCSC bigWigAverageOverBed utility, resulting in a mean phyloP score associated with each TE copy. TE phyloP scores were computed as the mean of all phyloP scores associated with all bases belonging to all copies.

#### DRS

Depletion Rank Scores computed by [23] were intersected with TE coordinates. Since DRS values are available as consecutive 500bp windows, the DRS value assigned to each TE copy was the average of the DRS of all windows overlapping the repeat weighted by the overlap length. The DRS score of a TE was then computed as the median of the DRS scores of its copies. Note that throughout the paper we use DRS’ = 1 - DRS/100 for sign consistency with the other two measures of selection (higher score corresponding to stronger conservation).

### Functional enrichment

Enrichments in GO:BP of TEs were obtained with the R implementation of GREAT [36] with default parameters. In the figures, we reduced the redundancy of the enrichment results using rrvgo [25] with default parameters, then we showed the top 10 terms sorted by P-value and then by fold enrichment. Enrichment P-values < 10^-300^ were set to 10^-300^. The enrichment results shown in the Supplementary data are the complete ones, before processing with rrvgo.

### Statistical analysis

For all linear models the dependent variable (i.e. the measure of phenotypic relevance) was inverse normal transformed and all independent variables were scaled to zero mean and unit standard deviation, so that the effect sizes are expressed in units of one standard deviation of the independent variable. All linear regression models were fitted with the *lm* function in R.

## Acknowledgements

Daniela Fusco is a PhD student enrolled in the National PhD in Artificial Intelligence, XXXVIII cycle, course on Health and Life Sciences, organized by Università Campus Bio-Medico di Roma. Davide Marnetto and Yari Cerruti were supported in part by the University of Turin through Grant For Internationalization 2022. The Genotype-Tissue Expression (GTEx) Project was supported by the Common Fund of the Office of the Director of the National Institutes of Health, and by NCI, NHGRI, NHLBI, NIDA, NIMH, and NINDS. The data used for the analyses described in this manuscript were obtained from the GTEx Portal on July 04 2022. We are grateful to Mathilde André, Elena Grassi, Giorgia Modenini, and Roberta Zeloni for insightful discussions.

## Data availability statement

The data underlying this article were accessed from the UCSC Genome Browser (https://genome.ucsc.edu/). The derived data generated in this research are available in the article and in its online supplementary material.

## Supplementary Tables

**Suppl. Tab. 1.** Regression coefficients and P-values of linear models predicting the density of eGenes from sequence features and measures of conservation. The adjusted R^2^ of the multivariable model is 0.44. UNI: univariable model. MULTI: multivariable model including all pairwise interactions.

| | $\beta_{\text{UNI}}$ | $P_{\text{UNI}}$ | $\beta_{\text{MULTI}}$ | $P_{\text{MULTI}}$ |
| --- | --- | --- | --- | --- |
| **DRS'** | 0.5438 | $7.117 \cdot 10^{-71}$ | 0.6105 | $2.371 \cdot 10^{-35}$ |
| **cCRE score** | 0.4416 | $1.488 \cdot 10^{-44}$ | 0.3324 | $5.714 \cdot 10^{-29}$ |
| **phyloP** | 0.1756 | $1.038 \cdot 10^{-7}$ | -0.3791 | $3.635 \cdot 10^{-6}$ |
| **phyloP:DRS'** | | | 0.227 | $5.495 \cdot 10^{-5}$ |
| **sequence age:DRS'** | | | 0.109 | $6.301 \cdot 10^{-5}$ |
| **cCRE score:sequence age** |  |  | -0.1651 | 0.000101 |
| **cCRE score:DRS'** |  |  | 0.07311 | 0.04715 |
| **sequence age:phyloP** |  |  | 0.09968 | 0.0477 |
| **sequence age** | 0.4598 | $1.325 \cdot 10^{-48}$ | 0.06159 | 0.1176 |
| **cCRE score:phyloP** |  |  | 0.04742 | 0.1374 |

**Suppl. Tab. 2.** Regression coefficients and P-values of linear models predicting the overlap with open chromatin from sequence features and measures of conservation. The adjusted R^2^ of the multivariable model is 0.45. UNI: univariable model. MULTI: multivariable model including all pairwise interactions.

| | $\beta_{\text{UNI}}$ | $P_{\text{UNI}}$ | $\beta_{\text{MULTI}}$ | $P_{\text{MULTI}}$ |
| --- | --- | --- | --- | --- |
| **cCRE score** | 0.5537 | $6.206 \cdot 10^{-74}$ | 0.5328 | $2.832 \cdot 10^{-66}$ |
| **DRS'** | 0.3518 | $8.576 \cdot 10^{-28}$ | 0.4105 | $6.974 \cdot 10^{-18}$ |
| **phyloP** | 0.207 | $3.14 \cdot 10^{-10}$ | -0.6982 | $2.11 \cdot 10^{-17}$ |
| **sequence age:phyloP** | | | 0.3543 | $2.3 \cdot 10^{-12}$ |
| **cCRE score:sequence age** | | | -0.2305 | $4.83 \cdot 10^{-08}$ |
| **phyloP:DRS'** |  |  | 0.2104 | 0.0001605 |
| **cCRE score:phyloP** |  |  | 0.09126 | 0.003972 |
| **sequence age** | 0.2312 | $1.828 \cdot 10^{-12}$ | -0.112 | 0.004142 |
| **sequence age:DRS'** |  |  | 0.06565 | 0.01471 |
| **cCRE score:DRS'** |  |  | -0.03304 | 0.3649 |

**Suppl. Tab. 3.** The 10 top TEs in terms of predicted effect on gene expression (“fitted” column). The clade of origin corresponds to the median sequence age of the TE copies (Methods). See also Suppl. Fig. 2.

| TE | fitted | repFamily | repClass | clade |
| --- | --- | --- | --- | --- |
| MamRep434 | 2.135 | TcMar-Tigger | DNA | eutherians |
| Tigger19a | 1.954 | TcMar-Tigger | DNA | eutherians |
| Mam_R4 | 1.887 | Dong-R4 | LINE | eutherians |
| MER94B | 1.789 | hAT-Blackjack | DNA | eutherians |
| hAT-N1_Mam | 1.781 | hAT-Tip100 | DNA | eutherians |
| Plat_L3 | 1.654 | CR1 | LINE | eutherians |
| AmnSINE1 | 1.523 | 5S-Deu-L2 | SINE | therians |
| Tigger16b | 1.513 | TcMar-Tigger | DNA | eutherians |
| CR1-3_Croc | 1.437 | CR1 | LINE | eutherians |
| MER57E3 | 1.38 | ERV1 | LTR | catarrhini |

**Suppl. Tab. 4.**
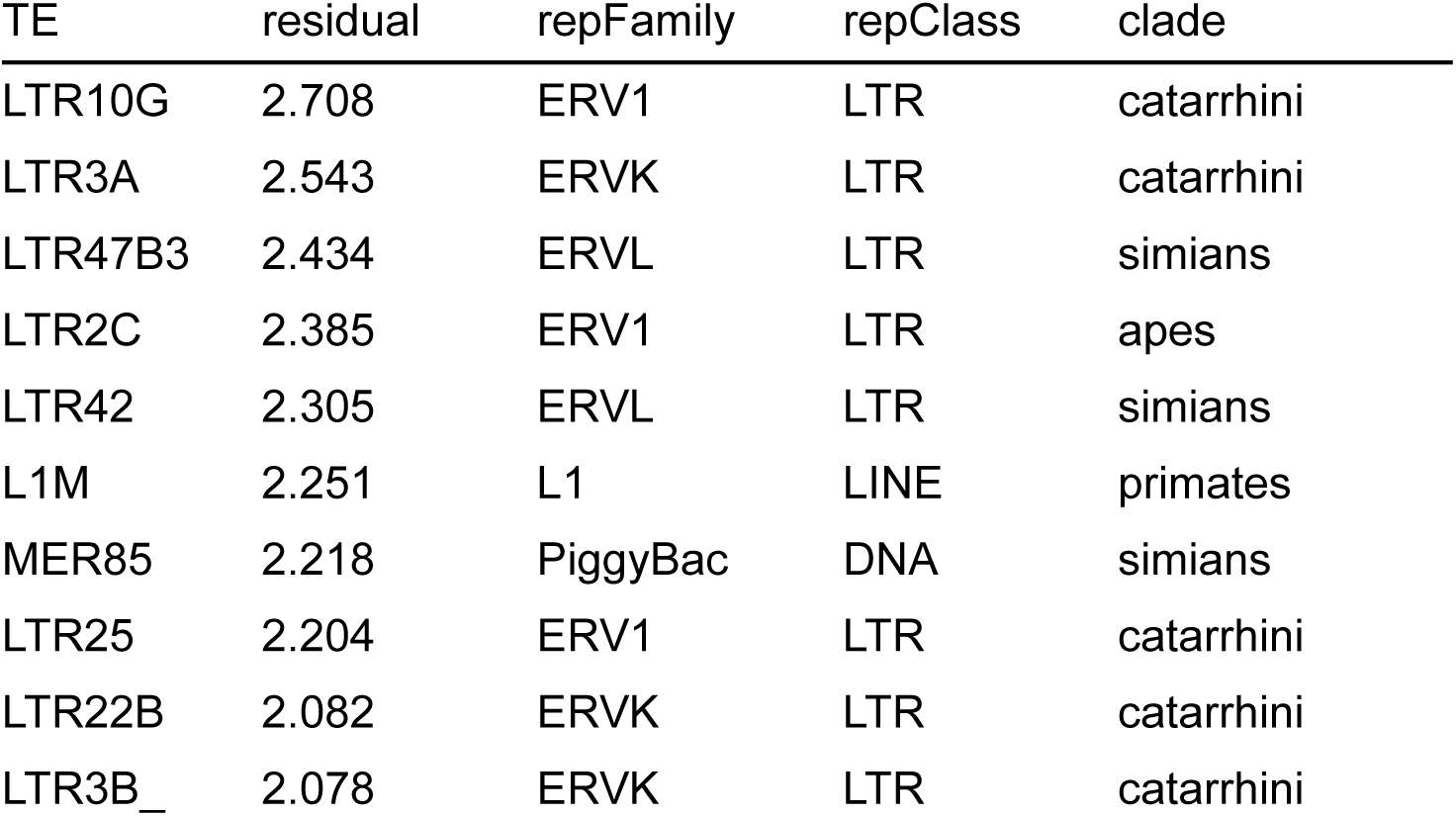
The 10 top TEs in terms of residuals of the model predicting the effect on gene expression. See also Suppl. Fig. 3.

**Suppl. Tab. 5.** The 10 top TEs in terms of predicted effect on chromatin accessibility (“fitted” column). See also Suppl. Fig. 4.

| TE | fitted | repFamily | repClass | clade |
| --- | --- | --- | --- | --- |
| MER121 | 2.961 | hAT | DNA | therians |
| AmnSINE1 | 2.187 | 5S-Deu-L2 | SINE | therians |
| MER126 | 2.066 | DNA | DNA | therians |
| MamRep434 | 1.795 | TcMar-Tigger | DNA | eutherians |
| Mam_R4 | 1.647 | Dong-R4 | LINE | eutherians |
| MER57E3 | 1.552 | ERV1 | LTR | catarrhini |
| Tigger19a | 1.448 | TcMar-Tigger | DNA | eutherians |
| Plat_L3 | 1.357 | CR1 | LINE | eutherians |
| LTR22C0 | 1.291 | ERVK | LTR | catarrhini |
| HERVK13-int | 1.253 | ERVK | LTR | african_apes |

**Suppl. Tab. 6.** The 10 top TEs in terms of residuals of the model predicting the effect on chromatin accessibility. See also Suppl. Fig. 5.

| TE | residual | repFamily | repClass | clade |
| --- | --- | --- | --- | --- |
| LTR19C | 2.863 | ERV1 | LTR | catarrhini |
| MER61F | 2.792 | ERV1 | LTR | simians |
| LTR2B | 2.368 | ERV1 | LTR | apes |
| LTR21A | 2.264 | ERV1 | LTR | catarrhini |
| LTR10B2 | 2.087 | ERV1 | LTR | catarrhini |
| LTR9C | 2.066 | ERV1 | LTR | simians |
| LTR3B_ | 2 | ERVK | LTR | catarrhini |
| LTR10A | 1.965 | ERV1 | LTR | catarrhini |
| LTR13 | 1.926 | ERVK | LTR | great_apes |
| LTR3A | 1.908 | ERVK | LTR | catarrhini |

## Supplementary Figures

**Suppl. Fig. 1.**
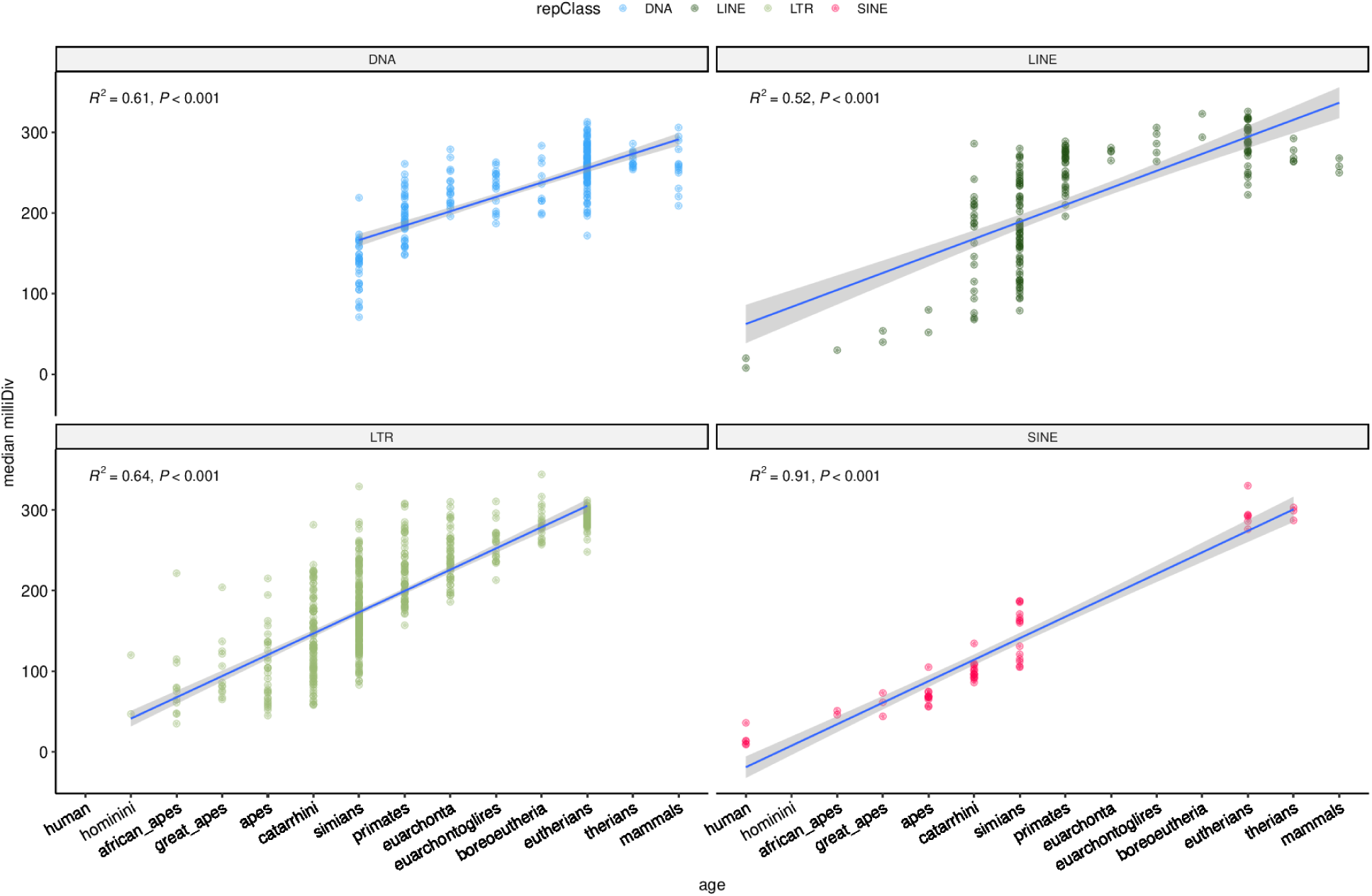
Comparison of TE age derived in [20] from vertebrate multiple alignments and in [16] from the divergence from the consensus sequence.

**Suppl. Fig. 2.**
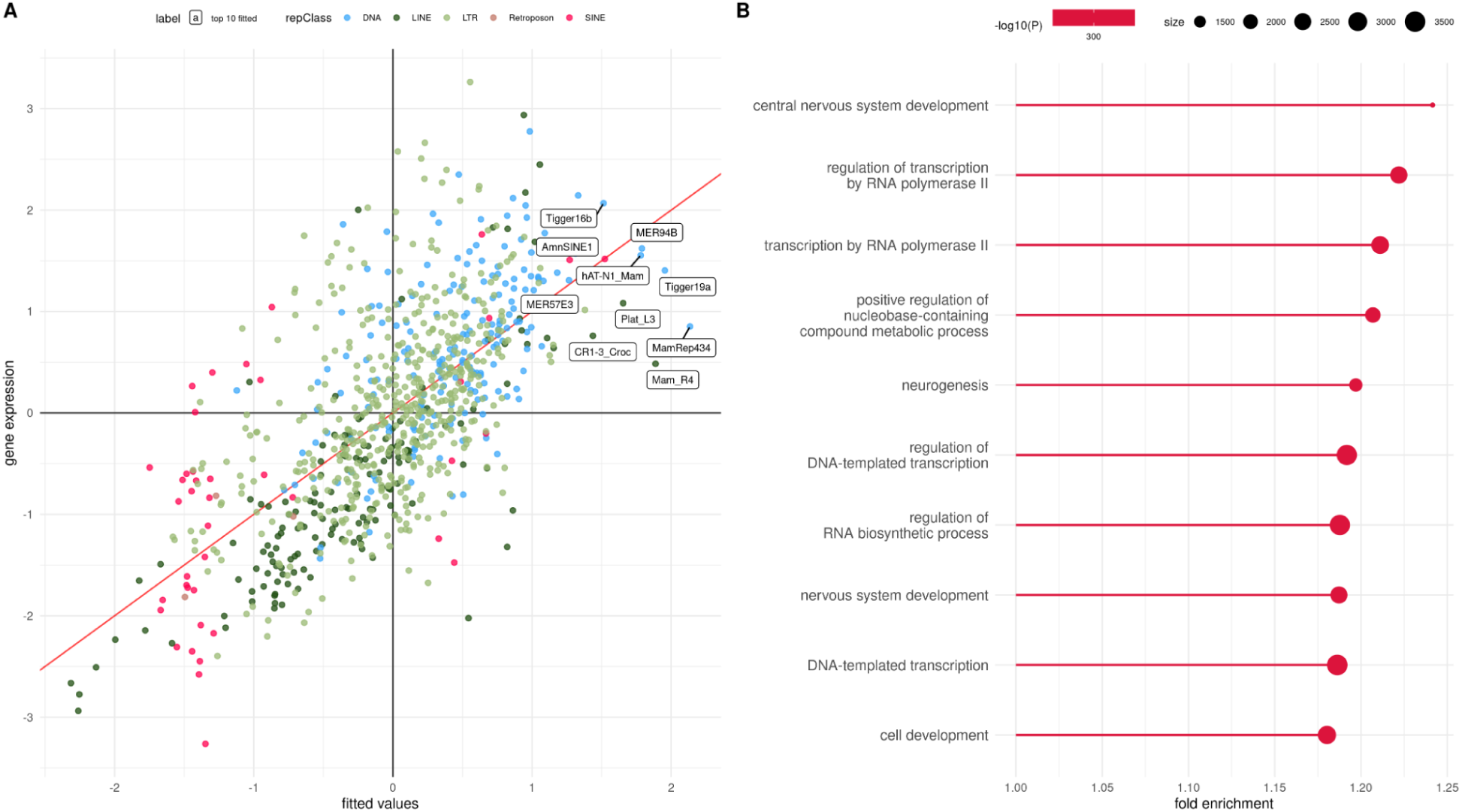
A: Linear model predictions (x axis) vs actual eGene density (after inverse normal transformation) of TEs. The labels indicate the top 10 TEs by predicted value. B: Gene Ontology terms enriched in the top 10 TEs by predicted value.

**Suppl. Fig. 3.**
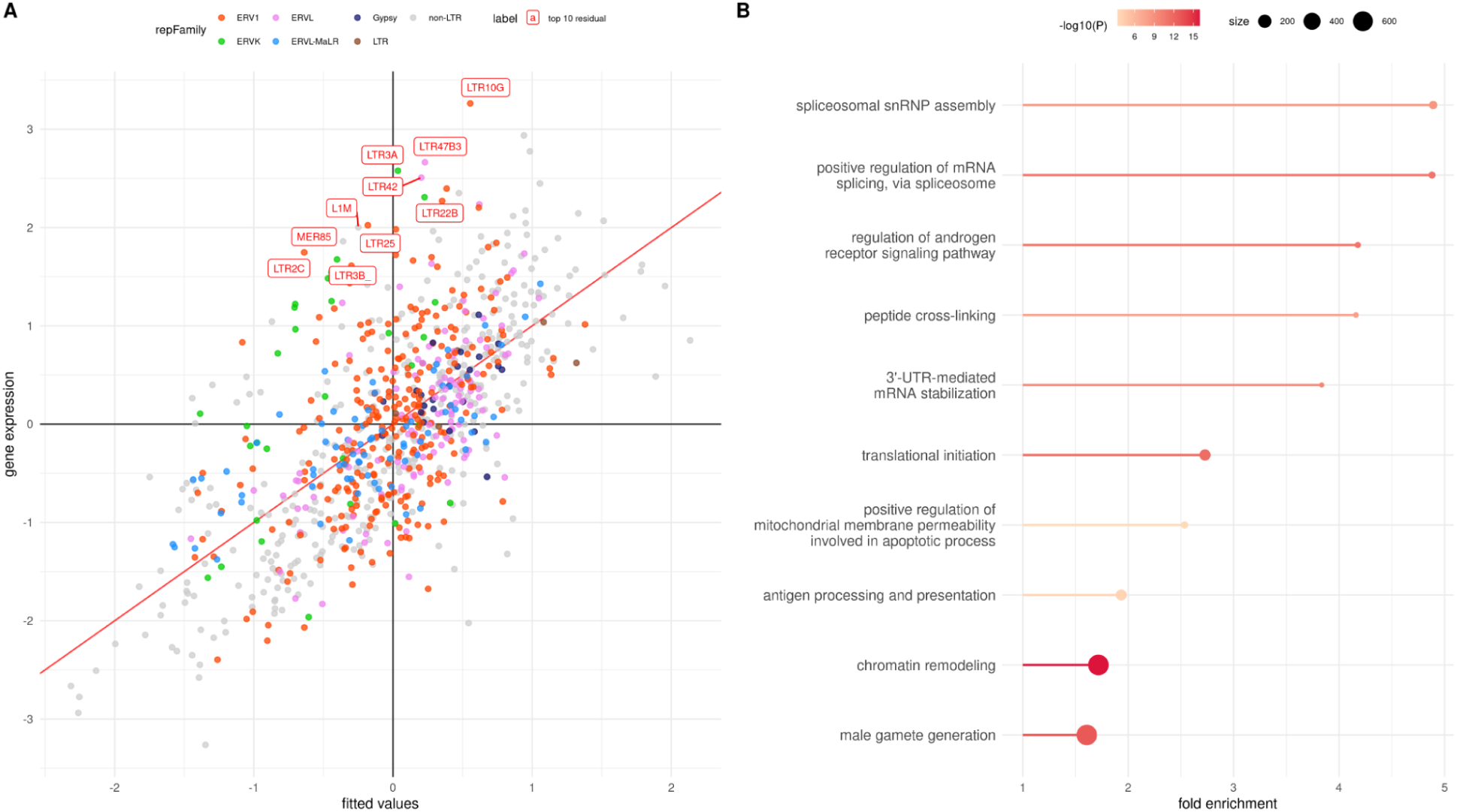
A: Same as Suppl. Fig. 2, except that the labels now indicate the top 10 TEs by model residual. B: Gene Ontology terms enriched in the top 10 TEs by model residual.

**Suppl. Fig. 4.**
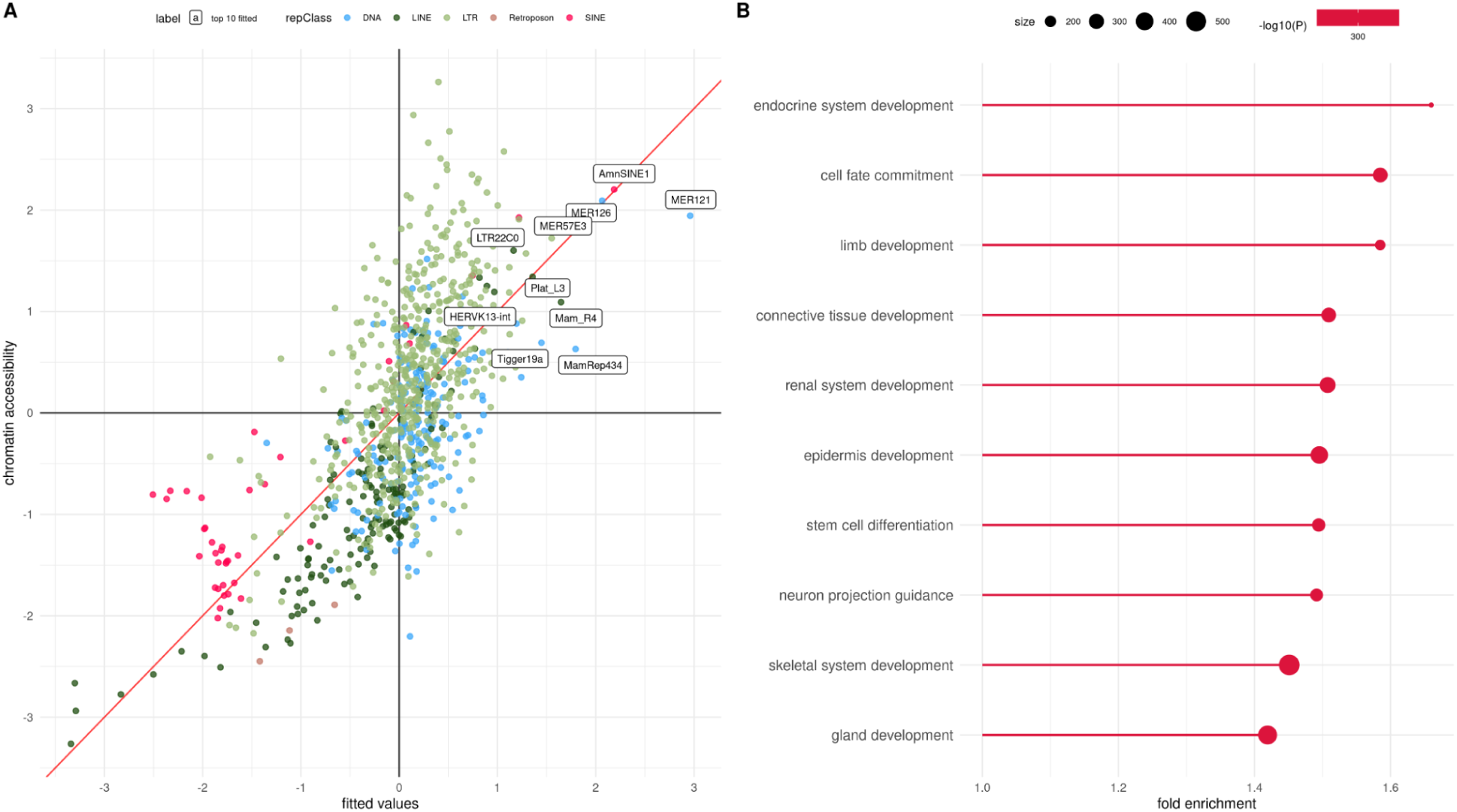
A: Linear model predictions (x axis) vs actual density of overlap with cCREs (after inverse normal transformation) of TEs. The labels indicate the top 10 TEs by predicted value. B: Gene Ontology terms enriched in top 10 TEs by predicted value.

**Suppl. Fig. 5.**
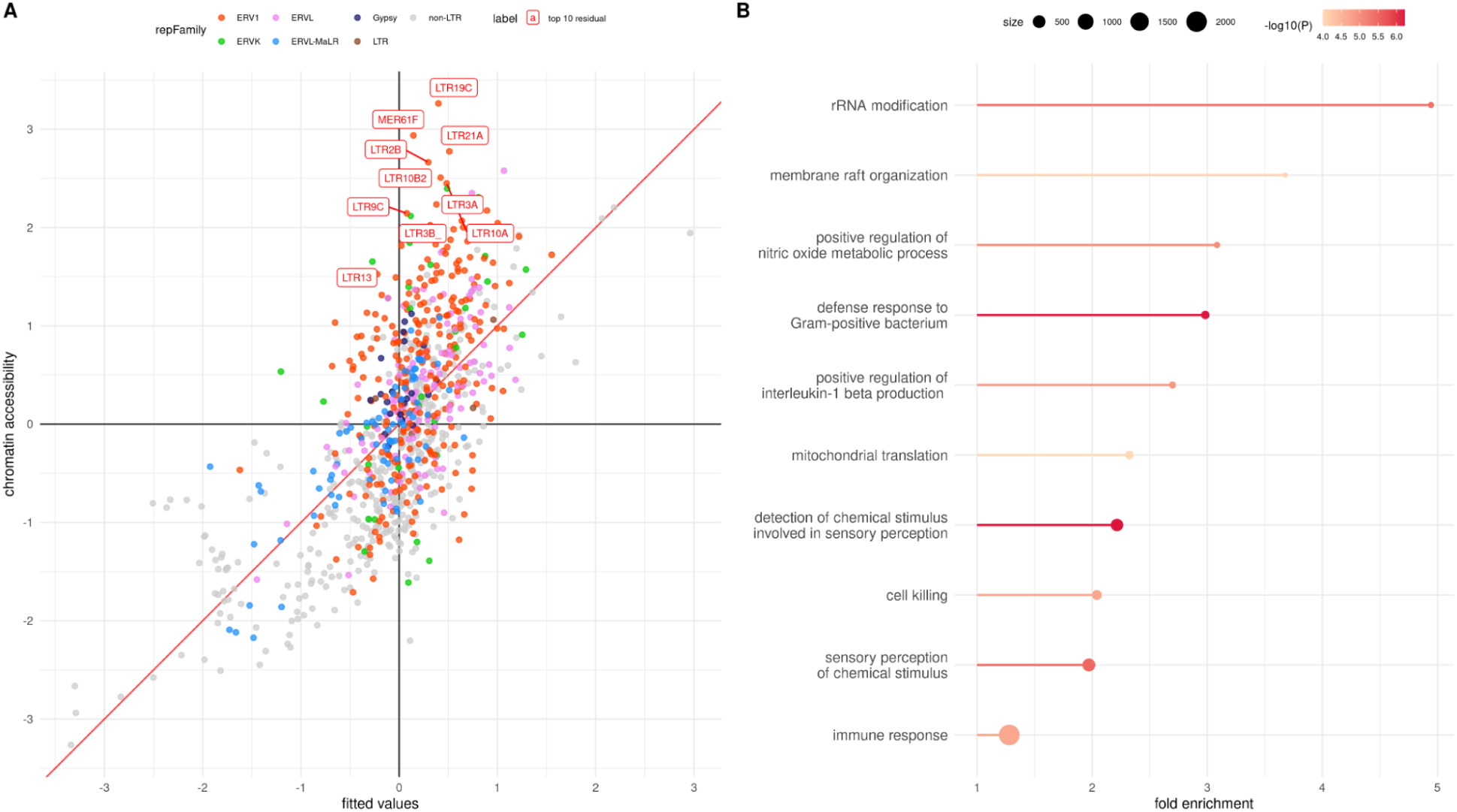
A: Same as Suppl. Fig. 4, except that the labels now indicate the top 10 TEs by model residual. B: Gene Ontology terms enriched in the top 10 TEs by model residual.

**Suppl. Fig. 6.**
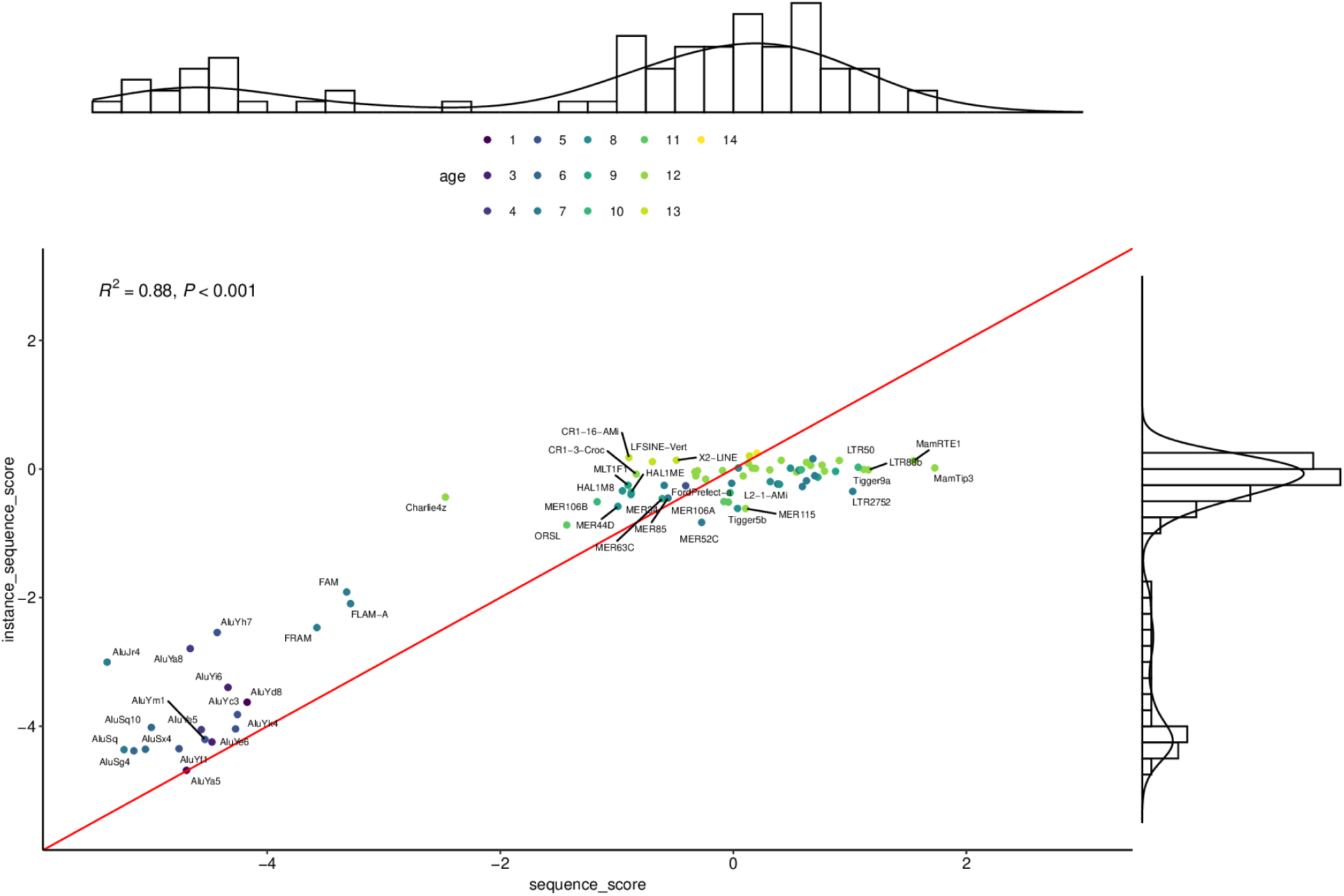
cCRE scores computed for 80 randomly chosen TEs using the consensus sequence (x axis) or averaging the score of all individual copies

